# A global survey of mycobacterial diversity in soil

**DOI:** 10.1101/562439

**Authors:** Corinne M. Walsh, Matthew J. Gebert, Manuel Delgado-Baquerizo, Fernando T. Maestre, Noah Fierer

## Abstract

*Mycobacterium* is a diverse bacterial genus ubiquitous in many soil and aquatic environments. Members of this genus have been associated with human and other animal diseases, including the nontuberculous mycobacteria (NTM), which are of growing relevance to public health worldwide. Although soils are often considered an important source of environmentally-acquired NTM infections, the biodiversity and ecological preferences of soil mycobacteria remain largely unexplored across contrasting climates and ecosystem types. Using a culture-independent approach by combining 16S rRNA marker gene sequencing with mycobacterial-specific hsp65 gene sequencing, we analyzed the diversity, distributions, and environmental preferences of soil-dwelling mycobacteria in 143 soil samples collected from across the globe. The surveyed soils harbored highly diverse mycobacterial communities that span the full-extent of the known mycobacterial phylogeny, with most soil mycobacteria belonging to previously undescribed lineages. While the genus *Mycobacterium* tended to have higher relative abundances in cool, wet, and acidic soil environments, several individual mycobacterial clades had contrasting environmental preferences. We identified the environmental preferences of many mycobacterial clades, including the clinically-relevant *M. avium* complex that was more commonly detected in wet and acidic soils. However, most of the soil mycobacteria detected were not closely related to known pathogens, calling into question previous assumptions about the general importance of soil as a source of NTM infections. Together this work provides novel insights into the diversity, distributions and ecological preferences of soil mycobacteria, and lays the foundation for future efforts to link mycobacterial phenotypes to their distributions.

**Importance:** Mycobacteria are common inhabitants of soil, and while most members of the bacterial genus are innocuous, some mycobacteria can cause environmentally-acquired infections of humans and other animals. Human infections from nontuberculous mycobacteria (NTM) are increasingly prevalent worldwide, and some areas appear to be ‘hotspots’ for NTM disease. While exposure to soil is frequently implicated as an important mode of NTM transmission, the diversity, distributions and ecological preferences of soil mycobacteria remain poorly understood. We analyzed 143 soils from across the globe and found that the genus *Mycobacterium* and lineages within the genus often exhibited predictable preferences for specific environmental conditions. Soils harbor large amounts of previously-undescribed mycobacterial diversity, and lineages that include known pathogens were rarely detected in soil. Together these findings suggest that soil is an unlikely source of many mycobacterial infections. The biogeographical patterns we documented lend insight into the ecology of this important group of soil-dwelling bacteria.

## INTRODUCTION

Mycobacteria are a diverse and ubiquitous group of Actinobacteria that have been well-studied given their potential importance as pathogens and their prevalence in a wide variety of habitats including many soil and aquatic environments (1). The best-studied members of the genus *Mycobacterium* include the obligate pathogens *M. tuberculosis* and *M. leprae*, but there are over 175 additional described members of the genus that are collectively referred to as non-tuberculous mycobacteria (NTM) (2). While many of these NTM taxa are innocuous, some are important pathogens of humans and animals (3, 4). NTM infections are predominantly acquired from the environment, and are increasingly recognized as a threat to public health worldwide (5). While the exact sources of these human NTM infections often remain unclear, water and soil are widely considered the predominant environmental reservoirs of NTM (1, 4, 6). A growing number of studies have focused on potentially pathogenic NTMs in aquatic systems, including residential plumbing systems (7–11), lakes (12, 13), and other aquatic environments (14–18). Much less is known, however, about the biodiversity and ecological preferences of NTM in soil and the relevance of soil-derived NTM to human health.

Members of the *Mycobacterium* genus are commonly detected across a broad range of soil types using both cultivation-dependent and cultivation-independent methods. We know from cultivation-independent surveys of soil bacterial communities that mycobacteria are nearly ubiquitous in soil and the genus *Mycobacterium* is consistently one of the more abundant genera of soil bacteria, ranging in relative abundance from ~0.5-3.0% of the total community (19–21). For instance, the *Mycobacterium* genus has been detected in soils from bogs (22), forests (13, 23, 24), croplands (25, 26) and livestock farms (27–29), and even in potting soil (30). However, much less is known about the ecological preferences of soil *Mycobacterium*. Current knowledge derived from a few studies suggests that mycobacteria are particularly abundant in high-latitude boreal forests (13, 24), with some research suggesting higher abundances in more acidic soils (17, 22, 25). However, studies identifying the specific ecological preferences of mycobacteria at both the genus and species or strain level of resolution across a range of soil and ecosystem types are lacking.

There are three main reasons why we lack a comprehensive understanding of mycobacterial diversity and distributions in soils across the globe. First, the majority of studies investigating environmental mycobacteria have relied on cultivation-based approaches to survey abundances and diversity (18, 31–33). While such approaches are clearly useful for addressing specific research questions, typical cultivation-based approaches are likely to underestimate the mycobacterial diversity found in environmental samples due to well-known culturing limitations (32, 34, 35). As such, it is likely that many environmental mycobacteria have remained uncharacterized (7, 36). Second, although PCR-based methods can be used to more broadly survey soil mycobacterial diversity (20, 27, 37, 38), most of these cultivation-independent studies have relied on the PCR amplification and sequencing of regions of the 16S rRNA gene, which often does not provide sufficient resolution to differentiate between distinct species or strains within the *Mycobacterium* genus (28, 37, 38). Alternate genetic markers such as the heat shock protein gene (hsp65) have proven useful for resolving closely-related mycobacterial species, but to date these marker gene sequencing approaches have been used predominantly for clinical isolate identification (39–42). Third, preexisting studies have generally focused on a relatively small number of soil samples with limited geospatial coverage. A more comprehensive understanding of the diversity and distributions of soil mycobacteria will clearly benefit from cultivation-independent analyses that provide greater taxonomic and phylogenetic resolution of those mycobacteria found across a broad range of soil types.

To advance our understanding of the ecology and biogeography of soil-dwelling mycobacteria, we analyzed a broad range of distinct soils collected from across the globe (143 locations) via cultivation-independent amplicon sequencing of two different marker genes (the 16S rRNA gene for genus-level analyses and the hsp65 gene for more detailed analyses of specific soil mycobacterial taxa). Using this global-scale survey, we addressed the following questions: 1) Which soils harbor the highest relative abundances of taxa within the *Mycobacterium* genus? 2) How does the diversity of mycobacterial communities vary across contrasting ecosystem types? and 3) What are the ecological preferences of individual mycobacterial lineages?

## RESULTS

### GLOBAL VARIATION IN RELATIVE ABUNDANCE OF GENUS MYCOBACTERIUM

We first used 16S rRNA gene sequencing to assess the genus-level relative abundances of *Mycobacterium* in each of 143 soil samples collected from across the globe (described in (21)). These soil samples were collected from a broad range of ecosystem types including tropical rainforests, arctic tundra, temperate grasslands, and hot deserts [Figure S1; Table S1]. We found that the *Mycobacterium* genus was widespread and reasonably abundant across the broad range of soil types examined here. 16S rRNA gene sequences from members of the *Mycobacterium* genus could be detected in all 143 samples, and this genus was consistently one of the more abundant named genera of bacteria identified. Out of 398 identifiable named genera, *Mycobacterium* was the 12^th^ most abundant genus across all soils by rank, ranging from 2^nd^ – 67^th^ across individual soils [Figure S2]. The relative abundance of the genus *Mycobacterium* was highly variable across the soils studied, ranging from 0.03% to 2.9% of all bacterial and archaeal 16S rRNA reads, with a median abundance of 0.52% [Figure S3].

The high degree of variation in the relative abundances of the genus *Mycobacterium* was, to some extent, predictable from the measured soil and site variables. We used Random Forest modeling (43) to identify the most important environmental predictors of the relative abundance of *Mycobacterium* across our samples. We found that ~25% of the variation in the relative abundance of *Mycobacterium* could be explained by the environmental variables measured. Temperature, aridity, and soil pH were the most important predictors of mycobacterial relative abundances [Table S3]. Mycobacteria were typically more abundant in soils collected from cold and temperate forests [Figure 1 D]. These patterns are consistent with the observation that mycobacteria generally had higher relative abundances in cooler, wetter and more acidic soils [FIG 1 A-C]. Across the 143 samples analyzed, mycobacterial relative abundances were, on average, higher in lower pH soils and in cooler climates [Fig 1 B, C]. Mycobacterial relative abundances were lower in drier (i.e. more arid) environments (44, 45) [Fig 1 A].

**FIGURE 1:**
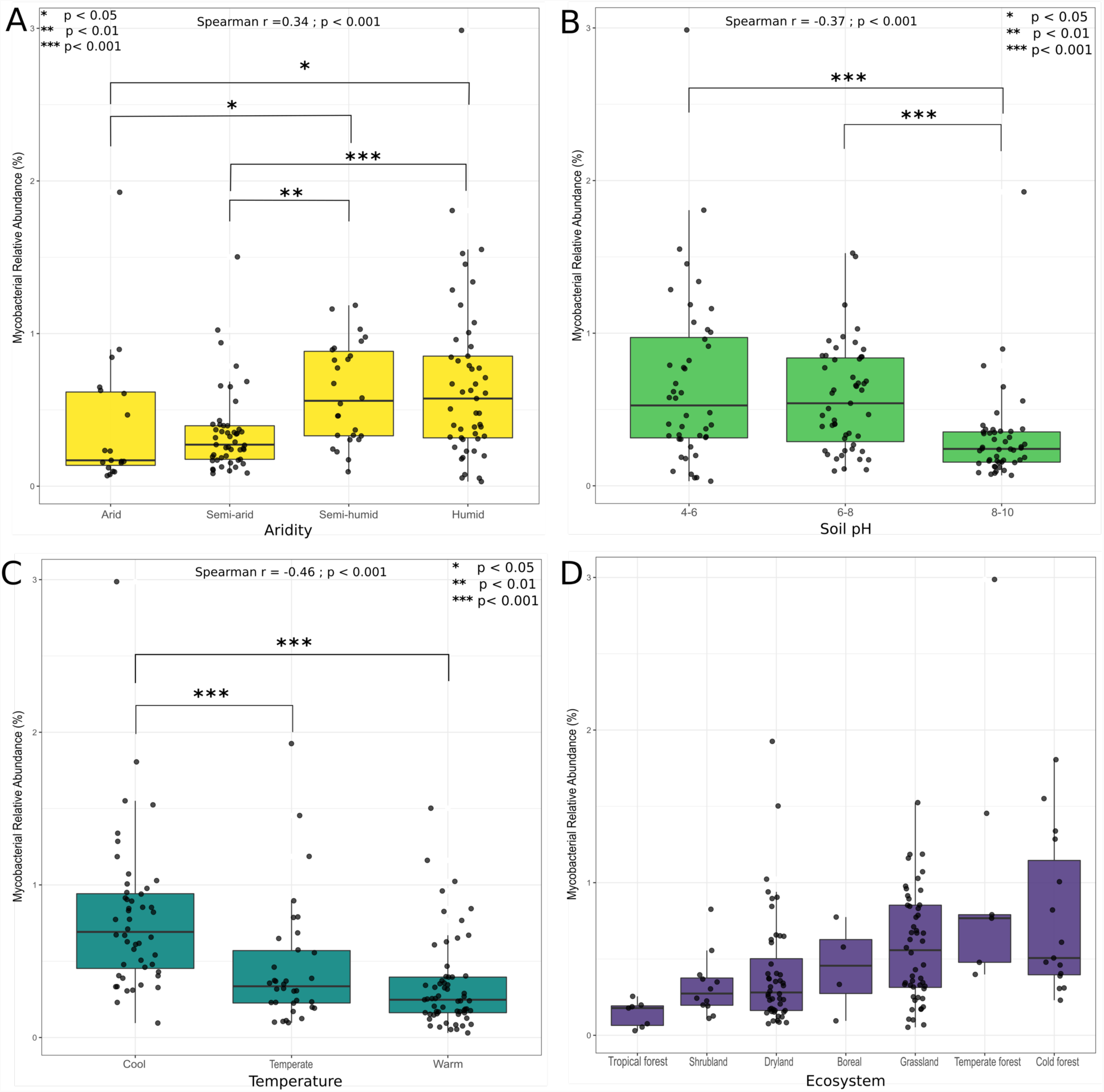
Relative abundances of mycobacteria across the 143 soils included in this study. Random forest analyses of 8 soil and site characteristics that described 25% of the variation in mycobacterial 16S rRNA gene relative abundances identified the most important environmental predictors as aridity index (A), soil pH (B), and minimum annual temperature (C). Boxplots show quartile ranges for each environmental category with black dots indicating exact sample values. Significant differences between groups (determined using pairwise Wilcoxon rank sum tests) are denoted by the asterisks with corresponding p-values. Spearman correlations suggest that mycobacteria are more abundant in wetter, more acidic, and cooler soils. Aridity categories were based on site aridity index values: “Arid” 0 - 0.2, “Semi-arid” 0.201 - 0.4, “Semi-humid” 0.401 - 0.7, “Humid” 0.701 - 2.5. Categories for Minimum Annual Temperature determined as: “Cool” - 35.0 to −9.0 °C, “Temperate” −9.01 to 1.0 °C, “Warm” 1.0 – 22.0 °C. Panel D shows differences in mycobacterial abundances across general ecosystem categories.

### GLOBAL DIVERSITY OF SOIL MYCOBACTERIA

The aforementioned 16S rRNA analyses do not allow us to comprehensively discriminate between different species or strains of mycobacteria. This is an important limitation given that the genus *Mycobacterium* contains a broad diversity of lineages with distinct phenotypic characteristics (46). Thus, we complemented the 16S rRNA gene analyses with amplicon sequencing of the hsp65 gene, which is widely used for phylogenetic analyses of the genus (47). These complementary analyses allowed us to identify specific lineages of mycobacteria and the distributions of these lineages across the 143 soils studied.

To streamline further analyses and investigate the ecological attributes of mycobacterial lineages, we used a phylogenetic approach to group closely-related mycobacterial sequences (exact sequence variants, or ESVs) into phylogenetic clades. Across all soils, the 472 mycobacterial ESVs that met our criteria for inclusion in downstream analyses (see Methods) fell into 159 distinct clades as identified by phylogenetic patristic distance. We clustered ESVs into phylogenetic clades because many sequences uncovered here were divergent from those in existing reference databases, creating challenges in confidently assigning taxonomic classifications.

Most of these mycobacterial clades were restricted in their distributions, and the number of different clades found in a single soil sample was highly variable. A median of 4 clades were identified per individual soil sample (range 0 to 18, mean 5.5, with 10 out of the 143 soils having no detectable mycobacterial hsp65 sequences). Most soils harbored only a few clades, while a few soils harbored a large number of clades [Figure S3]. No single clade was identified in all samples, and the most ubiquitous clade was found in 59 of the 143 soil samples. Together, these results highlight the high diversity of mycobacterial lineages that can be found in individual soils and that most soil mycobacteria lineages are restricted in their distributions.

Many well-characterized mycobacterial species, including known pathogens associated with human NTM diseases (including *M. abscessus M. chelonae M. fortuitum M. mucogenicum* (48, 49), were not detected in any of the 143 soils investigated here. In fact, the majority of soil-derived mycobacteria were so distantly related to described mycobacteria that they could not be assigned a species or strain identifier. When phylogenetically compared against a comprehensive mycobacterial database (47), only five of the 159 soil-derived clades clustered with known strains included in the Dai et. al. database (47) [Figure 2A]. Of the five clades that included a described isolate, only one clade included a species frequently implicated in disease [Figure 2B]. This pathogenic clade was comprised of members of the *M. avium* complex (MAC), which includes *M. avium, M. intracellulare*, and *M. chimera*. Representatives of the MAC clade were found in 15 of the 143 soil samples studied. The most ubiquitous and abundant clade identified across these soils comprised ESVs closely related to *M. novocastrense* and *M. rutilum*, found in 59 soil samples [Figure 2B]. We next assessed the geospatial distributions of the dominant mycobacterial clades and investigated the environmental conditions that best predict what clades are found in which soil types.

**FIGURE 2:**
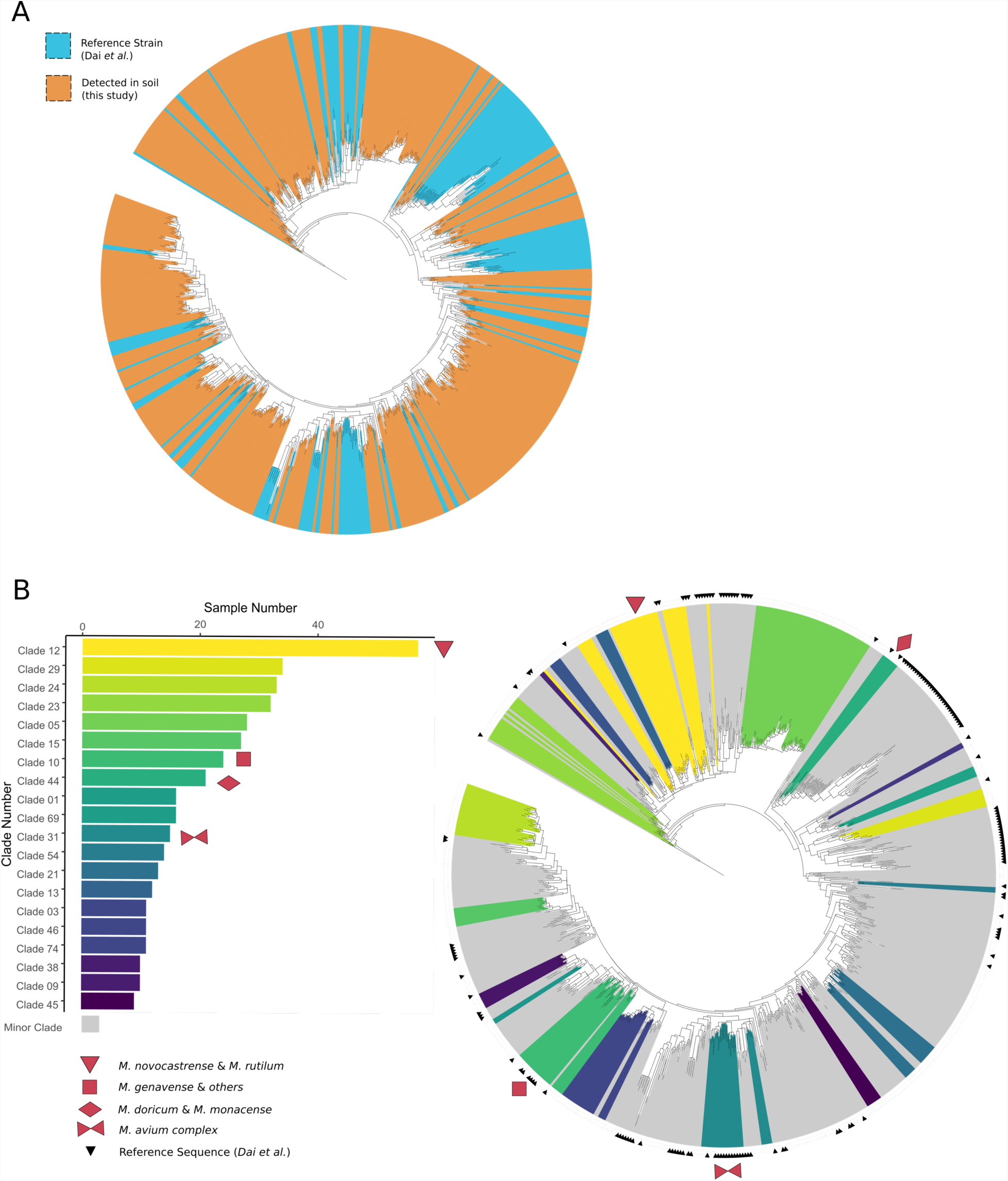
Phylogenetic tree representing described and previously undescribed mycobacterial hsp65 sequences. A) 472 Exact Sequence Variants (ESVs) were identified in soils sampled here, a majority of which represent novel and undescribed taxa. These ESVs span the known phylogenetic diversity of the genus. Colors indicate reference mycobacterial strains (blue) from Dai *et al.* (47) and sequences recovered from soils in this study (orange). B) Closely related ESVs were grouped into 159 clades based on patristic distance. The top 20 most ubiquitous clades are highlighted in color, with yellow colors indicating higher ubiquity (clades found in more soil samples). Red symbols indicate clades that include described taxa (indicated by the smaller black triangles). Both trees are rooted with the hsp65 sequence from *Nocardia farcinica* (DSM43665).

### BIOGEOGRAPHY OF THE DOMINANT MYCOBACTERIAL LINEAGES

Although most of the soil mycobacterial lineages identified were not closely related to described mycobacterial taxa, we explored their biogeographical distributions to gain insights into their ecologies. Likewise, by relating the presence of specific mycobacterial clades to soil or site characteristics we identified ‘hotspots’ of specific mycobacterial lineages that include potential pathogens. For these biogeographical analyses, we focused on the top 20 most ubiquitous clades (i.e. clades that were found in 10 or more soil samples), and used Random Forest modeling to identify the most important environmental predictors of clade presence or absence. Temperature, aridity, and soil pH were consistently the most important predictors of the distributions of ubiquitous mycobacterial clades [Table S3]. Distributions for 13 of the 20 clades could be predicted from the measured variables, with the majority of these predictable clades most likely to be found in soils that are acidic, cool, and wet. This finding agrees with the environmental preferences suggested by the genus-level 16S rRNA gene sequence data described above [Figure 1]. However, not all clades shared the same directional relationship with the measured environmental variables, highlighting the high levels of diversity and distinct environmental preferences within the mycobacterial genus [Figure 3, Table S4]. For example, the *M. rutilum* clade (clade 12) was more likely to be present in higher pH soils, while most other clades were more likely to be present in lower pH soils [Figure 3]. For a comprehensive list of all significant environmental predictors for each modeled clade see Supplementary Table 4. The *M. avium* complex (MAC clade) that includes known human pathogens was more likely to be found in wetter, more acidic soils [Figure S4]. While the number of samples containing members of this clade were too low to allow for robust modeling, Spearman correlations and Wilcoxon rank-sum tests support this apparent pattern (pH R = −0.41; Aridity Index R = 0.39).

**FIGURE 3:**
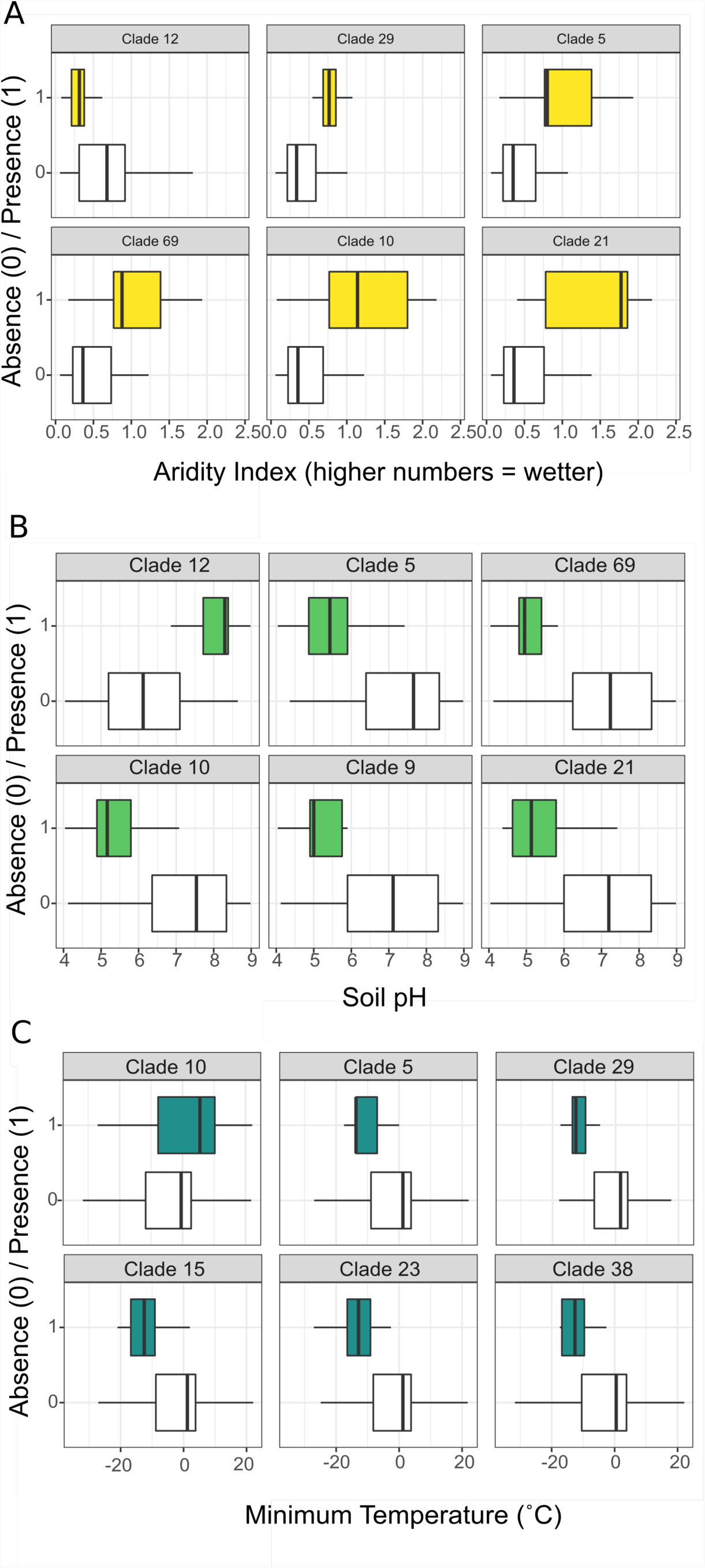
Environmental preferences of selected mycobacterial clades. The environmental conditions associated with presence of each clade shown here are significantly different from conditions in which the clade was absent, as determined by both Wilcoxon rank-sum tests and Spearman correlations (P < 0.05). Environmental variables that were consistently the most important for predicting clade presence in random forest models were aridity index (A), soil pH (B), and minimum annual temperature (C). Clades with notably strong environmental preferences are visualized in Figure 3, and a comprehensive list of all significant relationships with supporting statistics is provided in Table S4.

## DISCUSSION

The genus *Mycobacterium* is ubiquitous and reasonably abundant in soil as determined by our cultivation-independent 16S rRNA gene analyses. We found that *Mycobacterium* was typically one of the more abundant named genera of bacteria found in soil, confirming results reported in comparable studies (13, 19, 21). However, the relative abundance of mycobacteria was highly variable across soils ranging from 0.03 to 2.9% of 16S rRNA gene reads, a range similar to that reported previously (19–21). Some of this variation in total mycobacterial abundances could be explained from the measured soil and site characteristics, with mycobacterial relative abundances typically being higher in more acidic, colder, and moist soils. This observation is consistent with both cultivation-dependent and independent results published previously which have suggested that mycobacterial abundances tend to be higher in more acidic, wetter environments (16, 17, 22). However, we note that our model only explained 25% of the variance in genus-level mycobacterial abundances across the collected samples. Although such portion of explained variation is considered to be relatively high for comparable global-scale studies (50), unexplained variation could be a product of the fact that we did not measure all possible soil or site characteristics that can influence mycobacterial abundances, including the presence of amoebae (51, 52), specific organic carbon substrates (53), or the presence of livestock (25, 28, 29). Additionally, some of this unexplained variation could be a product of the coarse, genus-level analyses employed, which may not sufficiently capture the phenotypic differences or differences in niche preferences across taxa within the genus *Mycobacterium*.

As expected, soil mycobacteria were found to be highly diverse when analyzed with a higher resolution genetic marker (the hsp65 gene). Across all soils, we identified 472 ESVs that represented 159 distinct phylogenetic lineages with most of these lineages restricted to a small subset of soils (Figure 2). Not all mycobacteria were detected in all soils, an observation that could result from either dispersal limitations or, more likely, different mycobacterial strains having distinct environmental preferences. The latter explanation is supported by our results. In particular, we found that soil pH and climatic parameters were often the best predictors of the distributions of individual lineages (Figure 3), and that most of the lineages preferred low pH soils (a result consistent with the genus-level analyses, Figure 1). However, we also observed individual lineages that were more commonly detected in high pH soils (Figure 3). Likewise, while most lineages were mainly found in colder and wetter sites, others were more commonly detected in drier and warmer ecosystems (Figure 3). These results provide evidence that soil mycobacteria can exhibit contrasting, yet often predictable, environmental preferences.

Our findings further indicate that soils harbor a large amount of undescribed mycobacterial diversity. Of the 159 lineages detected in our global survey only 3% of the lineages (5/159) encompassed described isolates. These results contrast to a comparable survey of mycobacteria in household plumbing that used the same hsp65 marker gene sequencing approach, where the majority of the mycobacterial lineages included described taxa (7). The high proportion of undescribed mycobacterial lineages we recovered from soil is slightly higher than that reported previously in soils (20) and lakes (12) via 16S rRNA gene analyses. Of the small subset of soil mycobacterial clades that included described isolates, we found lineages that included *M. rutilum*, and *M. avium* complex, taxa that have previously been identified from cultivation-dependent analyses of soil mycobacterial communities (26, 53).

Only one of the detected lineages (*M. avium* complex) included potential human pathogens. Members of this group are frequently associated with clinical cases of NTM respiratory disease (17, 54). Interestingly, we found that the lineage including the *M. avium* complex (clade 31) showed strong preferences for wet and acidic soils, information that may be directly relevant to understanding the epidemiology of mycobacterial disease. However, as mycobacteria related to known pathogens were infrequently detected across the 143 samples studied, we think it is important to re-examine the widespread assumption that exposure to soil is an important mode of transmission of mycobacterial disease in humans. Specifically, outside of the *M. avium* complex, most of the mycobacterial isolates frequently recovered from patients with respiratory NTM disease such as *M. abscessus, M. mucogenicum, M. kansasii*, and *M. fortuitum* (3) were not detected in any of the soils. This differs from a similar investigation showing that mycobacteria frequently implicated in NTM disease to be common and relatively abundant in showerhead biofilms (7).

Our study represents one of the more comprehensive investigations of global soil mycobacterial diversity and the ecological factors shaping the distributions of individual soil mycobacterial lineages, many of which remain undescribed. We found that soil mycobacterial communities (both the *Mycobacterium* genus and individual lineages of mycobacteria) are often predictable from information on soil pH and site climatic conditions. Notably, most mycobacteria appear to prefer acidic, colder, and wetter soils, however, contrasting environmental preferences are found for individual clades. More generally, we show that mycobacteria can be abundant members of soil bacterial communities, but much of the soil mycobacterial diversity remains undescribed. Although the cultivation biases inherent in mycobacterial surveys are well-known (7, 36), our results suggest that this cultivation bias is particularly important in soil. We note that those mycobacteria commonly considered to be human pathogens are rarely detected in soils worldwide, challenging widely-held expectations that soil is an important source of NTM diseases. When we do detect lineages that include known pathogens, they are restricted in their distributions, information that is important for understanding the epidemiology of NTM respiratory disease and possibly other mycobacterial diseases of humans and livestock. Taken together, our study provides novel insights on the distribution and ecological preferences of soil mycobacteria in terrestrial ecosystems across the globe.

## METHODS

### Global soil sample collection

For detailed soil sample collection methods, see Delgado-Baquerizo *et al* (21). Here, we used a subset of 143 locations from Delgado-Baquerizo *et al* (21) for which soil was still available. Briefly, soils were collected from 143 locations across six continents spanning a wide range of ecosystem types (Table S1). Composite samples were collected to roughly 7.5cm depth, and were frozen immediately at −20°C. GPS coordinates and ecosystem type were recorded *in situ*, and climatic variables were determined using the Worldclim database (www.worldclim.org) (44, 55). Soil analyses were performed at the Universidad Rey Juan Carlos (Spain) following standardized protocols (56). Our database included information on key soil properties (organic carbon content, pH, C:N ratio, clay content, and electrical conductivity) and climatic variables (aridity index, annual minimum and maximum temperature) as explained in (21). Site aridity index is defined as precipitation/potential evapotranspiration (44, 45). For detailed information on soil and site characteristics, see Table S1.

### 16S rRNA gene sequencing to characterize the soil bacterial communities

We characterized the bacterial communities in our soils via 16S rRNA gene sequencing. For detailed sequencing methods, see Delgado-Baquerizo *et al* (21). Briefly, DNA was extracted from soils using the DNeasy Powersoil kit (Qiagen) following the manufacturer’s protocol. The V3-V4 hypervariable region of the bacterial 16SrRNA gene was amplified with 341F/805R primers, and sequenced on the Illumina MiSeq platform. Data were processed using the USEARCH10 pipeline (57). After quality filtering, reverse reads were trimmed before 10 bp and after 260 bp, merged (usearch8 -fastq_mergepairs). Following merging, 17 bp were trimmed off of the front of the merged reads to eliminate possibility of primer barcode contamination. Trimmed reads were filtered with a max error rate of 1.0, and a final truncation length at 410 (usearch10 -fastq_filter; 91.4% passed). Unique sequences were identified using usearch10 - fastx_uniques, and clustered at 97% identity to “Operational Taxonomic Units” (OTUs) with usearch10 -cluster_otus uniques.fa. Taxonomy was assigned with the Ribosomal Database Project Classifier(58) against the Greengenes 13_8 database(59), and chloroplast and mitochondrial reads were removed. To correct for differences in sequencing depth, samples were rarified to 10,000 reads per sample in R using the MCtoolsR package (https://github.com/leffj/mctoolsr/). The percentage of mycobacteria in each sample was calculated by summing reads classified to the *Mycobacterium* genus per sample.

### hsp65 gene sequencing to characterize mycobacterial diversity

To resolve mycobacterial taxonomic diversity beyond the genus level, we performed marker gene analysis of a region of the hsp65 gene specifically targeting mycobacteria. For detailed PCR and sequence methods, see Gebert *et al* (7). Briefly, a two-step PCR approach was used with mycobacterial-specific primers (60), and with the resulting amplicons (~400 bp) sequenced on the Illumina MiSeq platform. The sequence reads were processed using the uSEARCH pipeline (57). Raw reads were processed by trimming reverse reads to 250 bp (usearch8 - fastq_filter), merging (usearch8 -fastq_mergepairs), and quality filtering at a max error rate of 0.005 (usearch8 -fastq_filter -fastq_maxee_rate 0.005). Exact sequence variants (ESVs) were identified using uNOISE3 (61) (usearch10 -fastx_uniques and usearch10 -unoise3).

The hsp65 gene-targeting PCR primers do not exclusively amplify this marker gene from mycobacteria alone, as it can also amplify sequences representing other groups of actinobacteria such as *Nocardia* (62). To restrict our analyses to mycobacterial taxa alone, we filtered ESVs against a mycobacterial reference database (47), removing any reads that were <94% similar to the mycobacterial sequences in the reference database (usearch10 - usearch_global). Due to the large number of exact sequence variants remaining (2,836 mycobacterial ESVs out of 1,225,518 reads from all 143 samples), we only included those ESVs represented by at least 500 reads across all samples to focus on the more abundant and ubiquitous mycobacterial ESVs. After applying this threshold, 75% of all mycobacterial reads remained (926,258 reads, 472 ESVs).

### Phylogenetic tree construction and characterization of mycobacterial diversity

A majority of the soil-derived mycobacterial hsp65 sequences were highly divergent from reference sequences (see Results). Thus, to characterize the diversity of soil mycobacteria and for conducting downstream ecological analyses, we collapsed the ESVs into distinct phylogenetic clades. Phylogenetic relationships for the 472 ESVs were determined via maximum likelihood with RaxML (63). First, soil-derived ESVs were combined with the reference sequences from Dai et al.(47), and these sequences were aligned using MUSCLE v.3.8.31(64). Aligned reads were used to construct a tree with RaxML (raxmlHPC -f a -m GTRGAMMA -p 12345 -x 12345 -# 100), including *Nocardia farcinica* (DSM43665) as an outgroup for tree rooting. Sample and reference sequences were clustered into phylogenetic clades using RAMI (65), with a patristic distance threshold of 0.05 (5%). With the soil-derived ESVs combined with the reference sequences, this produced 242 clades containing 630 ESVs (472 mycobacterial sequences from our soils, 158 reference sequences). Many of the 242 clades included only reference sequences, as only 159 clades included soil-derived ESVs from this study. The tree was visualized and annotated using iTOL (66). Only five clades contained both reference sequences and soil-derived sequences. Sample sequences in these clades were confirmed to be relatives of the named taxa via BLAST (67), with all ESVs within a given clade sharing >99% identity with the corresponding reference strain. We determined the presence/absence of each clade in each sample, with ‘absence’ defined as a clade represented by less than 50 hsp65 reads in a given sample. Information regarding clade assignments for each ESV can be found in Supplementary Table 5.

### Statistical Analysis & Modeling

We used random forest analysis via the R package rfPermute (68) to identify the most important ecological predictors of genus-level mycobacterial abundance and mycobacterial clade distributions (functions rfPermute, rp.importance). Previous to these analyses, we pre-selected environmental variables from (21) that were not highly correlated (Pearson R < 0.7, Table S2). Based on these correlations and the relevance of remaining variables, the following environmental variables were included in downstream analyses: soil organic carbon content, pH, site aridity index, annual minimum and maximum temperature at the site, soil C:N ratio, soil clay content, and soil electrical conductivity. An independent random forest model was run separately for each clade of interest in addition to that built with the genus-level proportional abundance dataset (16S rRNA gene sequence data). Only models that described more than 20% of the variation were evaluated. “Important” environmental predictors in these models were defined as those that increased the Mean Standard Error of the model (MSE) more than 15% when excluded with p < 0.01.

To further understand the strength and directionality of the relationships between important environmental predictors and mycobacterial relative abundances or mycobacterial linage presence, we used Spearman rank correlations as we did not necessarily expect a linear relationship between the environmental variables and mycobacterial response. We then determined whether these correlations were associated with significant differences in mycobacterial relative abundances or clade presence for the environmental variables using the Wilcoxon rank sum test.

The statistical analyses to determine which soil or site characteristics were predictive of mycobacterial abundances (genus level analyses, 16S rRNA gene sequencing) or the presence/absence of individual mycobacterial clades (hsp65 gene sequencing) were performed in the R environment version 3.4.3 (69) with all plots generated using ggplot (70).

## ACKNOWLEDGEMENTS

We thank Hannah Holland-Moritz, Angela Oliverio, and Tess Brewer for their input and assistance in bioinformatics processing and statistical analyses. We also thank Victoria Ochoa and Beatriz Gozalo for their help with the laboratory analyses. Funding for this work was provided by grants to N.F. from the High Plains Intermountain Center for Agricultural Health & Safety and the Innovative Research Program of the Cooperative Institute for Research in Environmental Sciences, and to C.M.W. by the NSF IGERT Grant number 1144807 via the BioFrontiers Institute. The work of F.T.M. and the global drylands database were supported by the European Research Council [ERC Grant Agreements 242658 (BIOCOM) and 647038 (BIODESERT)].

